# The Ecosystem Pressure-Volume Curve

**DOI:** 10.1101/2022.09.12.507627

**Authors:** Oliver Binks, Patrick Meir, Maurizio Mencuccini

## Abstract

The ecosystem pressure-volume curve (EPV) is the relationship between vegetation water content and a representative value of water potential applied on a ground-area basis. The EPV attempts to reconcile our detailed and physically rigorous understanding of small-scale field-measureable processes to the spatial scale applicable to ecosystem and climate science. Successfully bridging that gap in scale potentially allows us to use field measurements to interpret remote sensing data, and then remote sensing data to inform our understanding of vegetation-climate interactions. Here we clearly define the idea of the EPV, evaluate the limitations of applying values of water content and water potential to ecosystems on a ground area basis, and discuss practical ways to construct the EPV with existing data. We also present the first EPVs based on data from nine different plots, including tropical rainforest, savanna, temperate forest, and a long-term drought experiment in Amazonian rainforest (Caxiuanã, State of Pará, Brazil). The initial findings suggest high levels of consistency among sites. In particular, the ratio of water to biomass across ecosystems appears to be constrained to around 1:3. Seven of nine sites had closely converging ‘relative maximum water storage’ (the proportion of total stored water than can be lost before an ecosystem succumbs to physiological damage) at 9.1% +/-1.8 standard deviation. Relative ecosystem capacitance may increase with site biomass (*P* = 0.091), but varied little across sites with a mean of 0.068 MPa^−1^ +/-0.029 standard deviation. These first estimates suggest that the EPV idea may reveal useful trends across ecosystems, potentially paving the way to increasing the ecophysiological significance of remote sensing data, and enabling an alternative method for modelling long-term ecosystem-climate feedbacks based on equilibrium thermodynamics.

## 1. Introduction

The relationship between water potential and water content in both plants and soil have long been of interest. Water potential (Ψ) is a thermodynamic property relating to the driving force for the movement of water in the environment. The relationship between Ψ and water content (θ) is known as the water release curve in soil and the pressure-volume curve in plants, and is dependent on the physical or biophysical properties of the soil or plant tissue in question. However, because of the scale at which land surface models operate, and the potential value of satellite remote sensing in providing ecosystem-scale information, there is mounting interest in understanding how these fundamental measures of water are linked at larger spatial scales (Konings *et al*., 2021). In this concept paper we explore whether Ψ and θ can be meaningfully scaled to represent an ecosystem, and if the relationship between these parameters at large-scale may offer a simple model of how ecosystems evolve to fill the hydraulic niche of the environment, and potentially respond to long-term environmental change.

Environmental moisture gradients, from wet to dry, are characterised by decreasing water content in the upper layers of soil (Guevara *et al*., 2021, Xie *et al*., 2015), increasing depth of the water table (Fan *et al*., 2013), decreasing minimum leaf water potentials (Peters *et al*., 2021, Sanchez-Martinez *et al*., 2020), and increasing evaporative demand (Zhang *et al*., 2017). A key component of evaporative demand is vapour pressure deficit (VPD), but atmospheric humidity can also be represented by water potential (Bannon, 2012, Kleidon, 2010, Thuburn, 2017). Therefore, gradients in aridity can be viewed as gradients in water potential of the soil, vegetation and atmosphere. Such gradients, from wet to dry, are additionally characterised by reductions in leaf area index (Iio *et al*., 2014, Yang *et al*., 2018) and biomass (Álvarez-Dávila *et al*., 2017, Stegen *et al*., 2011). Like biomass, the principal determinant of water contained in vegetation (vegetation water content, θ_ES_) is likely to be tree/forest volume (Calders *et al*., 2015, Vorster *et al*., 2020); therefore, one might hypothesise that large-scale gradients of ‘ecosystem water potential’ (Ψ_ES_) coincide with gradients of θ_ES_, resulting in a spatial equivalent of a pressure volume relationship.

Such a relationship may also be described in the context of temporal variations in Ψ_ES_ and θ_ES_. Transpiration causes steep vertical gradients in Ψ, making it challenging to conceptualise a single representative value of Ψ during periods of high flux rates. However, when transpiration is low (at night, or during periods of rainfall or drought) canopies tend towards values of Ψ_ES_ that are in equilibrium with the soil, e.g. predawn leaf water potential (discussed in Section 2), subsequently referred to as the equilibrium water potential^**1**^. In the medium-term (relating to seasonal time-scales and periods of drought from which an ecosystem can recover to pre-drought levels of function), one may expect variations in water availability to result in predictable changes in both Ψ_ES_ and θ_ES_. Long-term changes in water availability (i.e., multi-annual to century time scales) may cause ecosystem adjustments resulting in systems with lower biomass and leaf area index (Bennett *et al*., 2020, Brienen *et al*., 2015, da Costa *et al*., 2010). As in woody tissue (Oliva Carrasco *et al*., 2014, Rosner *et al*., 2019, Scholz *et al*., 2007, Tyree & Yang, 1990, Wolfe & Kursar, 2015), thresholds may exist at the level of the ecosystem that result in abrupt changes in the gradient of the PV relationship, providing avenues for ecosystem monitoring using remote sensing techniques.

Climate change is altering global patterns of evaporative demand, and will likely result in lower minimum atmospheric potentials (higher VPD (Barkhordarian *et al*., 2019, Ficklin & Novick, 2017, Yuan *et al*., 2019)), and shorter durations of maximum potentials, in much of the world. This will alter the fluxes, but may also alter ecosystem mean (or minimum) equilibrium water potentials. Given the predictable biogeographical patterns in global biome distribution (Holdridge, 1947, Humboldt, 1807), it is evident that plant communities tend towards a longer-term steady state determined by soil and atmospheric water potentials in terms of plant water potentials, biomass, and LAI etc. This being the case, an alternative approach to modelling fluxes, could be to model equilibrium states in response to longer-term average atmospheric conditions. In other words, atmospheric water potentials determine emergent structural characteristics of ecosystems, which in turn influence the average water content and water potential of the ecosystems, potentially providing a robust thermodynamic approach to modelling longer-term responses of ecosystems to gradual changes in climate. This would be a ‘state-based’ modelling approach (relating to the thermodynamic concept of a state function) as opposed to a process-based model. This approach, of measuring a state rather than a process, may also have utility in the context of longer-term and larger-scale environmental monitoring via remote sensing techniques.

This paper will describe the concept of the ecosystem water release curve and its possible applications, theoretical and practical. Water potential and water content tend to be treated differently in different media (soil, wood, leaves), therefore, the discussion will start with a description of each parameter in the environment in an effort to arrive at a consistent point of reference for the whole system. The concept of deriving a single value for ecosystem water content and potential will be reviewed in the context of deriving the theory, together with practical aspects pertaining to measurement in the field and use of existing data. We also present the first estimate of what ecosystem PV curves may look like, based on established theory and existing data.

## 2. Defining Ecosystem Pressure-Volume Terms

### 2.1. Summary of water potential in the environment

Chemical processes, including phase changes and diffusion, progress towards an equilibrium state in which gradients in chemical potential are fully dissipated. The hydrological cycle results from the continuous movement of water down a gradient of water potential towards an equilibrium state, and is perpetuated by the spatially and temporally variable input of energy across the Earth’s surface (Kleidon, 2010).

Following the pathway of water vertically upwards from its lowest point in a terrestrial system, we can define the water potential of the water table as 0 MPa, being free water at atmospheric pressure and assuming the osmotic potential is small. Above the water table, water is bound to the surface of soil particles and in pore spaces via capillarity, where the force of gravity, surface tension acting on menisci, and the resistance to the movement of water generates tension in the water column referred to as matric potential (negative hydrostatic pressure). The relationship between water content of the soil and Ψ is determined by the pore size distribution whereby larger pores empty initially at pressures closer to 0 Pa, while the smallest pores can retain water at substantially lower pressures.

In plants, the relationship is more complex where adjacent tissues can maintain Ψ equilibrium by balancing osmotic potential and hydrostatic pressure (Nobel, 1999, Pickard, 1981). In the xylem and in cell walls, pressure is the dominant determinant of water potential (referred to as tension and matric potential, respectively), and osmotic potential contributes minimally. In living tissues, water potential is determined by a combination of osmotic potential and turgor pressure.

The interface of the liquid-vapour phase change, in vegetation or soil, is typically the point of the system in which liquid water has its lowest chemical potential during evaporation. Evaporation and condensation are driven by the difference in chemical potential between the liquid and vapour. The evaporation of water reduces the hydrostatic pressure, thus Ψ, of the evaporative surface, and the resulting gradient in Ψ is transmitted through the vegetation and/or soil to the point at which Ψ is at its least negative value along the monotonic gradient of Ψ (Nobel, 1999).

The atmospheric potential oscillates diurnally according to temperature and humidity, typically achieving its lowest value (highest evaporative demand) around midday, and highest value at night during the formation of dew (when the gradient between the boundary layer and the liquid water surface is reversed) or during rainfall (Monteith & Unsworth, 2013). Consequently, the temporal mean water potential (daily, seasonal, annual etc.) at the evaporative sites, occurs as a function of the magnitude and timings (e.g. duration and frequency of extremes) of temporal cycles in atmospheric potential.

### 2.2. Defining ecosystem water potential, Ψ_ES_

Extreme horizontal gradients of Ψ at a given depth of soil should be uncommon due to the tendency of Ψ to equilibrate during periods of low flux, i.e., every night. Neighbouring plants, however, can differ in Ψ due to differences in physiological strategy and access to water (Sanchez-Martinez *et al*., 2020). Thus, horizontal spatial scaling of plant Ψ must be performed accounting for irregular spatial distributions, similar to the treatment of soil water *content* (Miller & Miller, 1956, Montzka *et al*., 2017, Warrick *et al*., 1977). Typically, there is a vertical gradient between the evaporative sites and the water table. Therefore, selecting a single value for Ψ_ES_ requires careful consideration of the representative position in the vertical column of the unsaturated zone.

We present three approaches for representing the vertical profile of water potential (i.e. the whole unsaturated zone) with a single value: i) weight Ψ according to water volume, ii) represent a relevant proportion of the vertical gradient, e.g. a mean of the upper two meters of the profile, or iii) simply the Ψ of the evaporative sites, i.e., leaves or the surface of the soil. These suggestions move from including the whole unsaturated zone to the smallest fraction of it, and have implications on the temporal dynamics of Ψ_ES_ which change more rapidly the closer to the evaporative sites and the smaller the volume of water that is being represented. However, because of the tendency for water potential to vertically equilibrate during periods of low evaporation, leaf predawn water potential essentially represents the volume of water between the canopy and the roots, accounting for the gravitational potential due to canopy height; hence, it is commonly used as a proxy for soil water potential.

Considering temporal scales, being a state-based, rather than a process-based, approach, Ψ_ES_ most usefully represents a steady-state value; that is, a longer-term average value arising due to the establishment of a stable gradient in water potentials between the atmosphere, vegetation and soil. However, this long-term value could be based on predawn (equilibrium) values, which represent the system at its highest potential (least stressed); midday values, representing the system at its lowest potential; or a combination of both. Practically, Ψ_ES_, can be determined by a combination of what is possible to measure in the field, and what is informative to models and remote sensing applications.

### 2.3. Summary of water content in the vegetation and soil

Water content, an extensive property (scales with volume), is conceptually simpler but can be expressed by different spatial dimensions depending on what is of interest. In soil it is standard to refer to θ in terms m^3^ m^−3^_soil_ (Hodnett & Tomasella, 2002, van Genuchten, 1980), in wood it can be expressed in kg m^−3^ (Meinzer *et al*., 2003, Scholz *et al*., 2007, Tyree & Ewers, 1991), while in leaves it is typically g m^−2^_leaf area_ or g kg^−1^_dry mass_. Additionally, water is often expressed as relative water content, which in soil is Θ_soil_ = (θ – θ_r_) / (θ_s_ - θ_r_) where θ_s_ is saturated water content (i.e. Ψ = 0 MPa), and θ_r_ is residual water content, a specified point at which dθ/dΨ tends to zero (van Genuchten, 1980). Wood and leaves are treated similarly but using mass, Θ_plant_ = (*M*_fresh mass_ – *M*_dry mass_) / (*M*_saturated mass_ – *M*_dry mass_), where again *M*_saturated mass_ is the mass of the leaf at Ψ = 0 MPa, but with the difference that *M*_dry mass_ is a state of zero water content (Wolfe, 2017). While absolute water content reflects the structure and makeup of the media, relative water content changes in reverse proportion to air content.

### 2.4. Defining ecosystem water content, θ_ES_

The scaling of water content thus presents the additional challenge that θ is a property of the medium, not the system. Therefore, θ does not tend to equilibrium with its surroundings, and adjacent media (e.g. leaves, wood, soil) may maintain very different water contents (absolute and relative), even at water potential equilibrium.

The notional horizontally averaged water content, θ_ES_, in the middle of a vertical profile of forest would contain sections of tree stem, leaves and large amounts of air. One could assume that the area of a slice, in order to converge on a horizontally averaged water content in a forest, would be in the order of hundreds to thousands of m^2^ depending on the structure of the forest. It becomes useful then to express water content per ground area, kg m^−2^ which is equivalent to mm thickness of water. Because the vast proportion of water in the unsaturated zone would be close to the water table, it would also make sense to choose a value that represents only the upper canopy and twigs. However, volume estimates of the upper canopy are likely to be very approximate, even when derived from lidar (Calders *et al*., 2015). Thus, a practical solution could be to treat θ_ES_ as the ground-area-based water content of the entire above ground biomass (AGB); thereby also enabling the use of existing biomass estimates.

AGB consists of four components that are likely to have different PV relationships: sapwood, heartwood, bark and leaves (see SI Section 1. Water stored in above ground biomass). Heartwood comprises 40 – 60% of wood volume in mature trees (Cermák *et al*., 2007, Knapic *et al*., 2006, Pérez Cordero & Kanninen, 2003, van der Sande *et al*., 2015), but the consensus is that it does not contribute substantially to plant hydraulic function (SI 1.2) (Holbrook, 1995). From a practical perspective, there is very little data on the contribution of heartwood water to transpiration, or long-term water balance, on which robust estimates could be based. Water in bark and leaves may contribute significantly to daily transpiration flux, but in absolute terms are likely to contain little water compared to the volume of water stored in stems. Thus, over a longer time scale, the total contribution of leaves and bark to changes in θ_ES_ is likely to minimal in most ecosystems (SI 1.3, 1.4, Table S1). Most experiments on wood water content and pressure-volume curves have been performed on sapwood, and it is reasonable to assume that sapwood water content varies most directly in response to canopy water potentials. Therefore, providing one can estimate sapwood as a fraction of biomass (SI 1.1.1), sapwood water content may be a useful means of characterising θ_ES_.

## 3. Generating ecosystem pressure-volume curves

We constructed the first ecosystem-level PV relationships with respect to medium- to longer-term changes in θ_ES_ and Ψ_ES_ using plot-level forest inventory data from the sites listed in Table 1. Here we define the ‘normal’ physiological range of plant water potential as being between the maximum possible canopy water potential with respect to tree height, and the measured minimum dry season canopy water potential (See SI Section 2. Choosing a threshold water potential). We based the analysis on the following assumptions, shown graphically in **figure 1**:

**Table 1.** Description of sites including mean annual temperature (MAT) and mean annual precipitation (MAP) and the sources of data.

| Site | Vegetation type | MAT (°C) | MAP (mm) | References |
| --- | --- | --- | --- | --- |
| Caxiuanã, Eastern Amazon, Brazil | Tropical rainforest | 23.0 | 2272 | (Bittencourt *et al.*, 2020, Chave *et al.*, 2014, Fisher *et al.*, 2007) |
| Litchfield, Australia | Wet tropical savanna | 27.8 | 1714 | (Peters *et al.*, 2021)<br>tern.org.au* |
| Tumbarumba, Australia | Temperate woodland | 9.8 | 1417 | tern.org.au* |
| Cow Bay, Australia | Tropical rainforest | 23.7 | 3700 | Binks *et al.* in review (Bradford *et al.*, 2014, Chave *et al.*, 2014) |
| Robson Creek, Australia | Tropical rainforest | 18.8 | 2264 | Binks *et al.* in review (Bradford *et al.*, 2014, Chave *et al.*, 2014) |
| Alice Mulga, Australia | Semi-arid savanna | 22.4 | 308 | (Peters *et al.*, 2021) |
| Great Western Woodlands, Australia | Temperate woodland | 18.5 | 268 | (Peters *et al.*, 2021) |
| Cumberland Plain, Australia | Temperate woodland | 17.7 | 809 | (Peters *et al.*, 2021) |

\*Terrestrial Ecosystem Research Network

**Figure 1.**
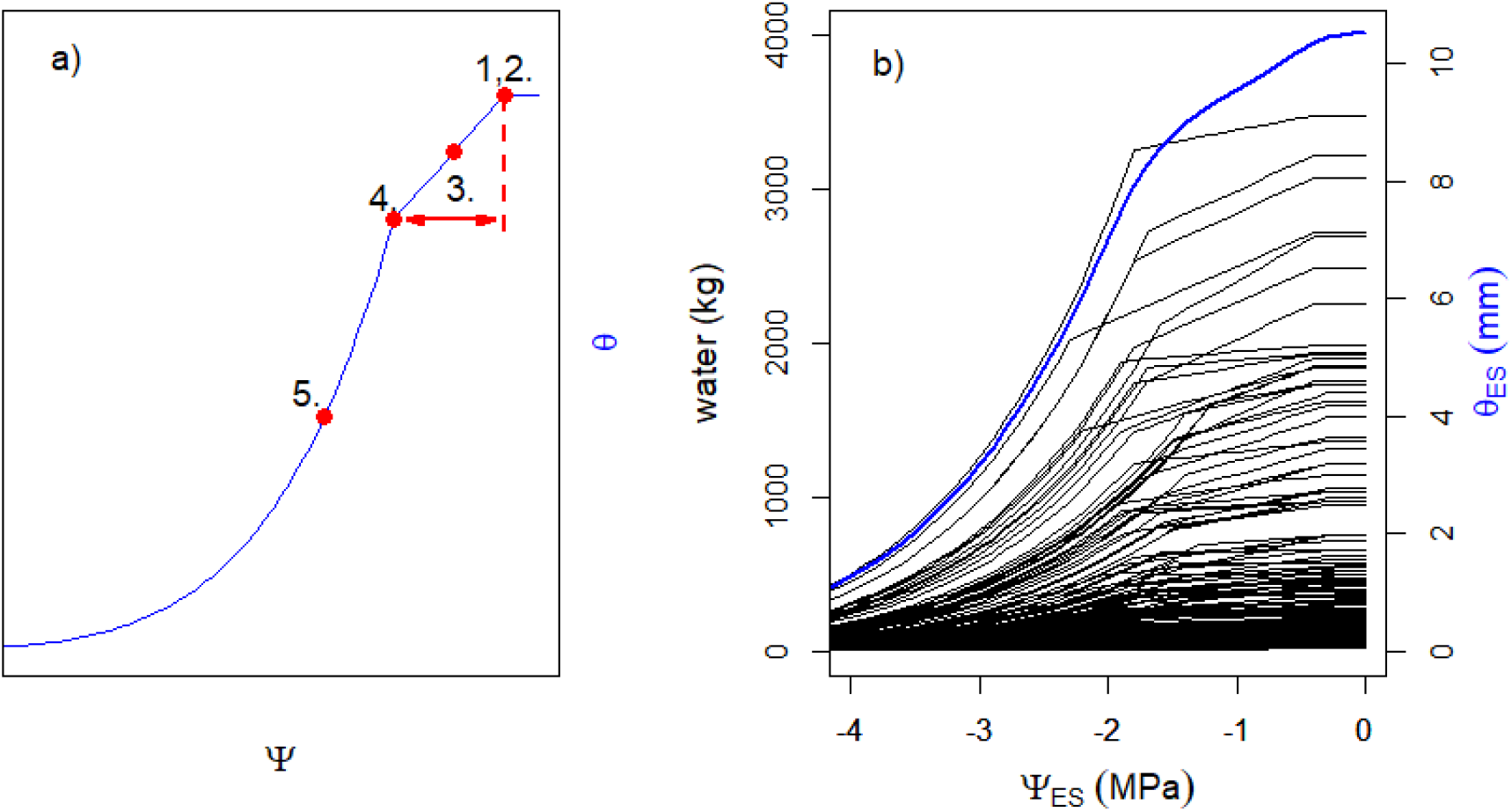
Panel a) shows a schematic relationship between water content (θ) and water potential (Ψ), i.e., a pressure-volume (PV) relationship, generated using the following parameters: 1. Saturated water content, Θ_sat_; 2. Maximum water potential, Ψ_max_; 3. Constant hydraulic capacitance throughout ‘normal’ physiological range indicated by the red double ended arrow, C_1_; 4. Threshold water potential at which the PV relationship transitions into non-linear region, Ψ_threshold_; 5. Exponentially declining capacitance as a function of water potential, C_2_(Ψ). In panel b) the black lines show modelled relationships between the absolute water content contained in the sapwood of individual trees (kg water) versus their equilibrium water potential, of all trees within a simulated one hectare stand. The blue line in panel b) represents the total ‘thickness’ of water in a forest (θ_ES_), as the sum of the water from all trees, with respect to the ‘ecosystem water potential’ (Ψ_ES_), blue line.

- Water potential is at equilibrium, meaning that the vertical gradient is minimal.
- The hydraulic capacitance within the ‘normal’ physiological range of water potentials, *C*_1_, is constant resulting in a linear relationship between Ψ and θ within that range (Meinzer *et al*., 2003, Oliva Carrasco *et al*., 2014, Scholz *et al*., 2007, Tyree & Ewers, 1991, Wolfe & Kursar, 2015, Ziemińska *et al*., 2020).
- Water content declines exponentially at Ψ values more negative than the normal range, consistent with available data showing pressure volume relationships of sapwood (see citations in point above), leading to a dynamic second phase capacitance, C_2_(Ψ).
- The most relevant pool of water varying over the medium-term (seasonal and drought events) occurs in the sapwood (Cermák *et al*., 2007, Holbrook, 1995), for reasons both physiological (i.e. declines in sapwood water status are directly related to mortality), and quantitative.

Rather than applying a single PV curve to the total amount of sapwood in a plot, a PV curve was generated per tree, and plot-level values were calculated from the combined properties of all trees (Fig. 1b). This approach avoids scaling errors associated with the Jensen’s Inequality (Ruel & Ayres, 1999); but it also enables the incorporation of random variability of parameters between individuals and species; and allows the addition of individual- or species-specific traits (e.g. wood density) resulting in different values of Ψ and θ(Ψ).

Scaling water relations traits to absolute values of water required an estimate for the volume of sapwood per tree: *V*_sw_tree_ = *F*_sw_·*AGB*_tree_/ρ = *F*_sw_·*V*_tree_, where *AGB*_tree_ is the above ground biomass of a single tree, ρ is wood density, and *F*_sw_ is the volume fraction of sapwood. See Table 2 for the descriptions, equations and parameter values for each of the variables, and SI 1.1.1 for derivation of *F*_sw_.

**Table 2.** Parameters used to derive the pressure volume curve for a single tree.

| Parameter<br>(units) | Description | Derivation |
| --- | --- | --- |
| $\Psi_{\max}$<br>(MPa) | Maximum (least negative) water potential | $H_{\text{tree}} \times -0.01$, where $H_{\text{tree}}$ is tree height (m), and -0.01 is a constant describing the gravitation effect on pressure ($\text{MPa m}^{-1}$) in a water column. |
| $\Psi_{\text{predawn}}$<br>(MPa) | Measured predawn canopy water potentials | See Table 1 for sources of data. |
| $\Psi_{\text{threshold}}$<br>(MPa) | Minimum safe water potential | This value is based on dry season midday leaf water potentials for the purpose of this analysis (See SI Section 2. Choosing a threshold water potential). Each tree was randomly allocated a value for $\Psi_{\text{threshold}}$ from a random normal distribution generated from the mean and standard deviation of midday leaf water potential values taken at a given site. |
| $\Theta_{\text{saturated}}$<br>($\text{kg m}^{-3}$) | Saturated sapwood water content | An empirical relationship reported by Dlouha et al. (2018) where $\Theta_{\text{saturated}} = -0.67 \cdot \rho + 1$, and $\rho$ is wood density. |
| $\theta_{\text{ES}}$<br>(mm) | Ecosystem water content | The sum of the sapwood water content of each tree in a plot expressed per ground area for a given equilibrium water potential. See equation 1 in the main text. |
| $C_1$<br>($\text{kg m}^{-3} \text{MPa}^{-1}$) | Sapwood capacitance of the linear phase of the pressure-volume curve. | Where field data exist, each tree was randomly allocated a value for $C_1$ from a random normal distribution generated from the mean and standard deviation of capacitance values. In the absence of field data, the mean value was derived from an empirical equation from Zieminska et al. (2020) of the form: $C_1 = -157.8 \cdot \rho + 137.7$, where $\rho$ is wood density, and the standard deviation was taken as $0.5C_1$. |
| $C_2(\Psi)$<br>($\text{kg m}^{-3} \text{MPa}^{-1}$) | Post-threshold capacitance of individual trees | Based on an empirical relationship designed to approximate the steep decline in water content with water potential following the threshold water potential e.g. (Christoffersen et al., 2016, Meinzer et al., 2003, Scholz et al., 2007, Wolfe & Kursar, 2015). $C_2(\Psi) = (-10 - \Psi_i)^n$, where '-10' is the minimum predicted water potential in this analysis, $i$ is the $\Psi$ at the $i$th step, and $n$ is a random number between and including 2 to 8 to reflect differences in $C_2(\Psi)$ between individual trees. |
| $C_{\text{ES}}$<br>($\text{mm MPa}^{-1}$) | Ecosystem hydraulic capacitance | The sum of the tree-level products of capacitance and sapwood volume over the plot area. See equation 2 in the main text. |
| $\rho$<br>($\text{g cm}^{-3}$) | Wood density | Used to derive $C_1$, $\Theta_{\text{saturated}}$, and $V_{\text{AGB}}$ |
| AGB<br>($\text{kg ha}^{-1}$) | Above ground biomass | Taken from existing datasets |
| $V_{\text{AGB}}$<br>($\text{m}^3$) | Volume of above ground biomass | Derived by dividing AGB by average wood density: $V_{\text{AGB}} = \text{AGB}/\rho$ |
| $F_{\text{sw}}$<br>(dimensionless fraction) | Sapwood as a fraction of total volume | $F_{\text{sw}} = 2.9 \cdot \text{DBH}^{-0.6}$, derived as a compromise between the empirical relationship presented by Cordero & Kanninen (2003), and the ratio of sapwood area to basal area from Kunert et al. (2017), Moore et al. (2018), Aparecido et al. (2016) and Wang et al. (2010). See SI 1.1.1 for full details on deriving $F_{\text{sw}}$. |
| $H_{\text{tree}}$<br>(m) | Tree height | Available in the datasets. |

The tree-level PV curves were generated using only five fundamental parameters (Fig. 1). Water potentials were simulated from Ψ_min_ to 0, designated Ψ_i_, where Ψ_min_ = -10 MPa. Water content for Ψ_i_ ≥ Ψ_threshold_, was calculated as θ_tree_(Ψ_i_) = Θ_sw_sat_ – C_1_(Ψ_max_ – Ψ_i_). At Ψ_i_ < Ψ_threshold_ the capacitance transitions into the non-linear zone changing as a function of Ψ, C_2_(Ψ_i_) = (Ψ_min_ – Ψ_i_)^*n*^, and θ_tree_(Ψ_i_) = C_2_(Ψ_i_)·θ_tree_(Ψ_threshold_). Refer to Table 2 for derivation of C_2_

The threshold water potential, Ψ_threshold_, was based on dry season midday water potentials (Ψ_md_), which represents a low Ψ from which plants can rapidly recover full hydraulic function (SI Section 2). Critically, Ψ_threshold_ represents a point at which the system is at water potential equilibrium, while the canopy is at Ψ_md_. Thus, Ψ_md_ = Ψ_threshold_ when Ψ_md_ = Ψ_predawn_. Individual trees were allocated a value for Ψ_threshold_ taken from a random normal distribution based on the mean and standard error of reported Ψ_md_ (Table 2).

Ecosystem water content for each water potential, θ_ES_(Ψ_i_), was calculated as the sum of the sapwood water content of all trees per plot (*n*), at a given water potential, expressed by ground area of the plot (*A*_plot_)

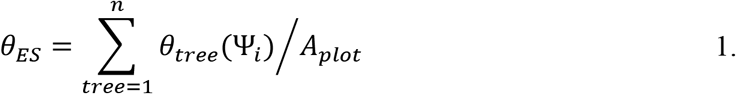

Ecosystem capacitance (*C*_ES_) was calculated as:

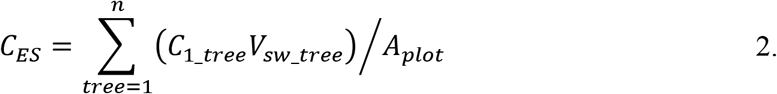

Where *C*_1_tree_ and V_sw_tree_ are the first phase capacitance and sapwood volume of each tree in a plot, respectively.

‘Accessible water’ was taken to be the difference in θ_ES_ between Ψ_predawn_ and a low water potential from which full plot-level physiological recovery was possible. The lower boundary was taken to be Ψ_threshold_ + 1 standard deviation (of measured Ψ_md_) in order to account for the distribution of Ψ_threshold_ of the individual trees – half of which would be less negative than plot mean Ψ_threshold_. It should be further emphasised that this does not represent the difference in predawn and midday θ_ES_, because the Ψ values here are taken as equilibrium values i.e., without steep vertical gradients in Ψ.

‘Maximum storage’ was calculated as the difference in θ_ES_ between Ψ_max_ (the maximum possible water potential of the canopy, given canopy height and the assumption that all water comes from the soil), and Ψ_threshold_ + 1SD (as above). This represents the biggest possible seasonal change in θ_ES_ without the occurrence of physiological damage.

## 4. Results and discussion

The purpose of expressing pressure volume curves of ecosystems (Fig. 2) is to bridge the gap between physiologically relevant variables of the kind collected in field studies with the scale at which ecological processes influence regional and global climate, and can be detected using emerging remote sensing technology. We speculate that there is a relationship between biomass, water content and climate, which links physiology with large-scale and long-term ecosystem characteristics.

**Figure 2.**
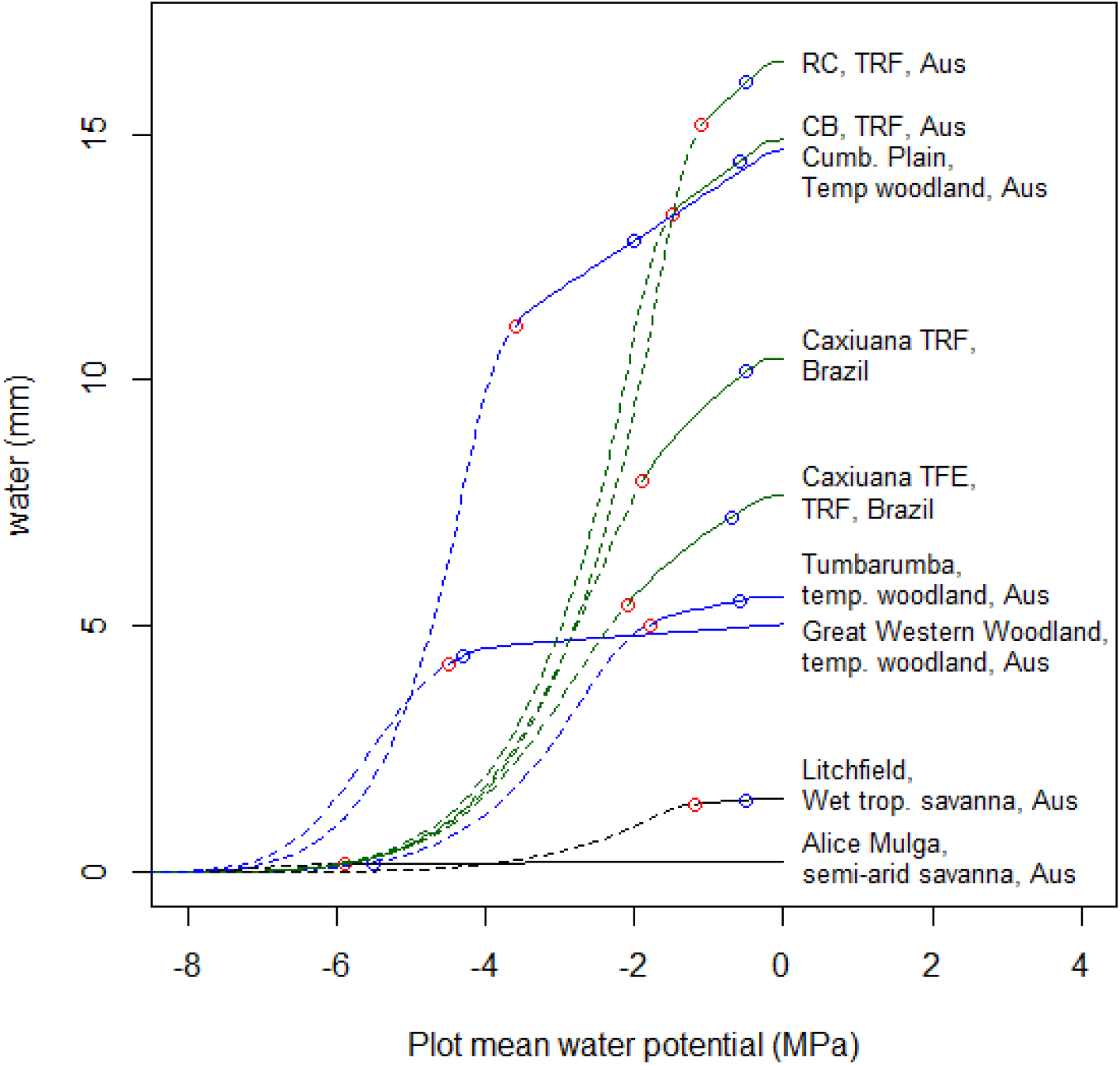
Ground area based water content versus equilibrium water potential of various ecosystems. Labels on the figure refer to the sites, where TRF stands for tropical rainforest, and ‘Caxiuanã TFE’ refers to a through-fall exclusion drought experiment in the Eastern Amazon where through-fall has been reduced by approximately 50% since 2002. Solid lines are based on data, while the dashed lines represent approximations of the pressure-volume relationship following the water potential threshold. Tropical rainforest systems in green, temperate forest in blue, and tropical savanna in black. The points on each curve represent equilibrium water potentials as measured at predawn (blue) and the threshold (red) water potentials.

Water potential in plants is highly physiologically constrained. Midday atmospheric water potentials (Ψ_atm_) above forest canopies are typically between -50 and -100 MPa, while water potential in the upper surface of forest soil can also be very low at less than -10 MPa (Binks, unpublished data). Yet, there is little data to suggest that plant canopies in any ecosystem routinely sustain water potentials less than -5 MPa (e.g. Fig. S4). Maintaining high water potentials requires investment of carbon (e.g. into roots and vascular tissue resilient to cavitation), but the fundamental trade-off between carbon uptake and water loss, means that as evaporative demand increases, all else equal, carbon assimilation must decrease. We propose that the carbon-water trade-off in fluxes amounts to the same trade-off in stores, resulting in a robust, mechanistically-driven dependence between biomass and ecosystem water content, but at a longer timescale than is usually considered when describing ecological mechanisms. The implication is that biomass can be viewed as an ecosystem-level functional trait emerging from the physiological requirement of vegetation to maintain water potentials within a tolerable range. This view is supported by the preliminary evidence presented here showing a constant ratio between water content and above ground biomass across ecosystems including semi-arid savannah, temperate woodland and tropical rainforest (Figures 3 and 4).

**Figure 3.**
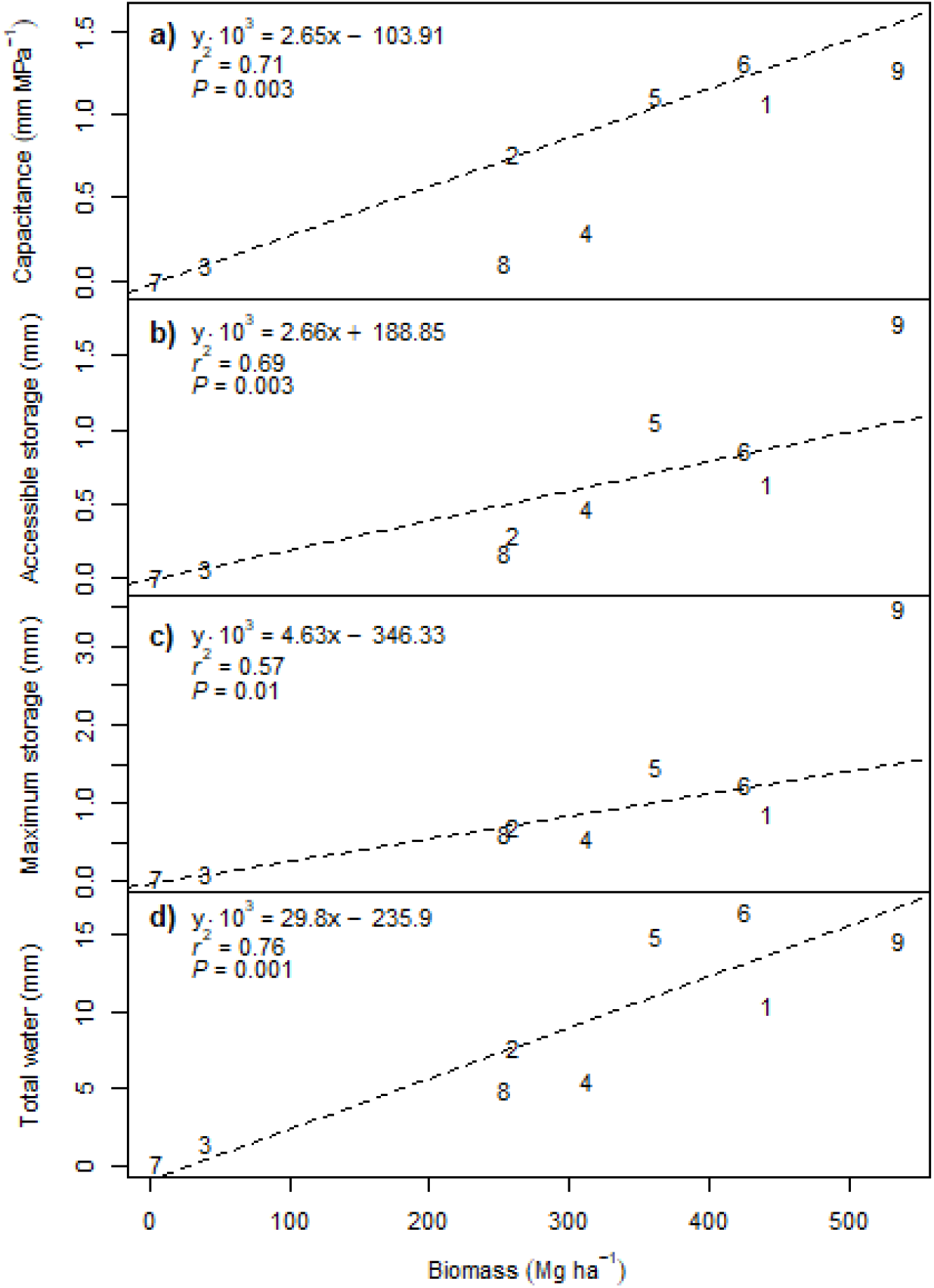
Plot-level hydraulic capacitance (a), accessible stored water between measured predawn water potential and the threshold water potential; (b), water storage between maximum hydration and threshold water potential (c), and the total amount of water stored in vegetation at saturation (d), all in relation to plot biomass. Each number on the plot represents data from each site described in Table 1 corresponding to: 1. Caxiuanã (non-droughted TRF); 2. Caxiuanã (droughted TRF); 3. Litchfield (wet trop. Savanna); 4. Tumbarumba (temp. woodland); 5. Cow Bay (TRF); 6. Robson Creek (TRF); 7. Alice Mulga (semi-arid savannah); 8. Great Western Woodland (temp. woodland); 9. Cumberland Plain (temp. woodland). Coefficients from linear regression are multiplied by 10^3^ to reduce the number of significant figures.

**Figure 4.**
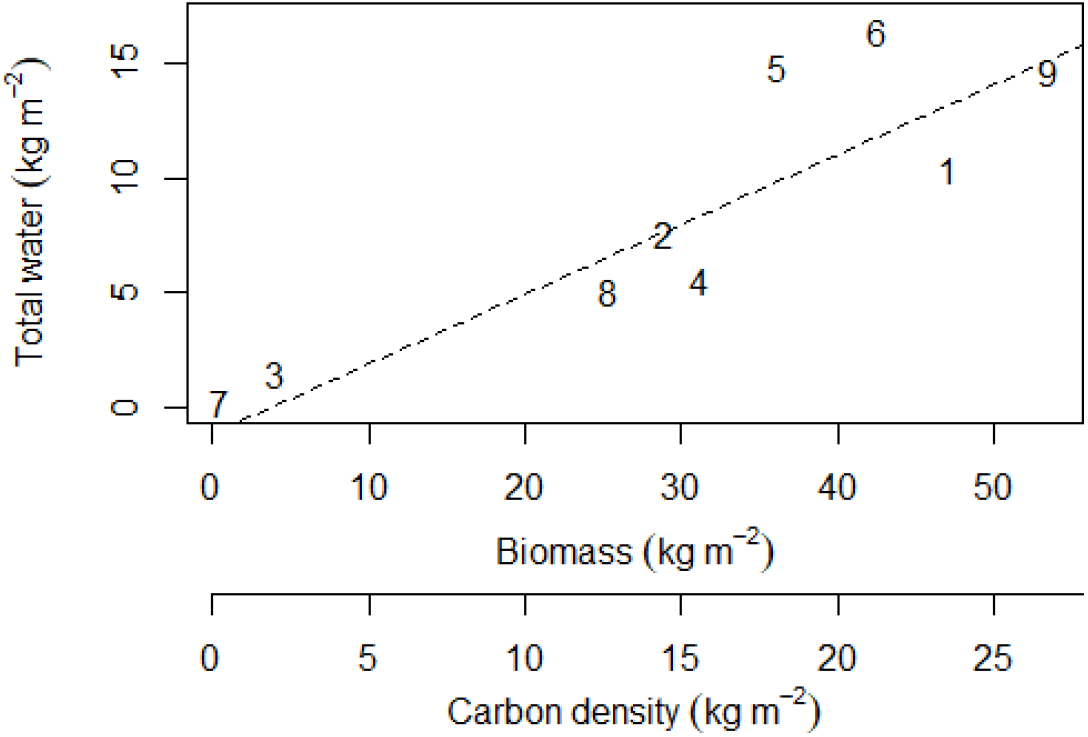
Comparison of the total amount of water stored in vegetation at saturation with total biomass and carbon density (biomass/2) expressed in the same units. The coefficients are of course similar to those expressed in Fig. 3d, where Total water = 0.30*Biomass = 0.59*Carbon density. Each number on the plot represents data from each site described in Table 1 corresponding to: 1. Caxiuanã (non-droughted TRF); 2. Caxiuanã (droughted TRF); 3. Litchfield (wet trop. Savanna); 4. Tumbarumba (temp. woodland); 5. Cow Bay (TRF); 6. Robson Creek (TRF); 7. Alice Mulga (semi-arid savannah); 8. Great Western Woodland (temp. woodland); 9. Cumberland Plain (temp. woodland).

### 4.1. Modelling applications

Vegetation has limited control over evapotranspiration at community level because of non-zero cuticular conductance of plant tissues and soil evaporation (da Costa *et al*., 2018). Therefore, modelling the water contained in ecosystems over long time-scales may be compared to modelling water levels in a lake, where changes in supply and atmospheric demand (the hydraulic environment) result in differences in ecosystem water content. Given that there appears to be a universal cross-biome ratio between water and above ground biomass, here estimated as approximately 1:3 water:biomass (Fig. 4), it is likely that long-term changes in the hydraulic environment would alter the total amount of stored water in an ecosystem causing predictable changes in biomass as the ratio becomes re-established.

As the purpose of this opinion paper is to introduce the concept of an ecosystem-level pressure-volume relationship, we will not present an analysis of the physical links between climate, ecosystem steady-states of water content and potential, and biomass, here. Nevertheless, one can envisage how long term reductions in water can reduce the volume of living tissue in an ecosystem (e.g. (da Costa *et al*., 2010, Meir *et al*., 2015)) and, conversely, how ecological pressures may act to fill a hydrological niche, maximising biomass within the constraints of other limiting resources, where/when water limitation becomes reduced.

### 4.2. Remote sensing applications

Ideally, remote sensing applications will act to monitor water stress within ecosystems, and help us to identify tipping points where vegetation may transition from one type to another (Konings *et al*., 2021). The water contained within the ecosystems represented here can vary over the medium term (seasonal-annual time scales) by a maximum of around 10% of the maximum water content (Figures 5 and 6c) without incurring physiological decline irrespective of ecosystem biomass or typical midday water potentials, based on the assumptions in the analysis (Martinez-Vilalta *et al*., 2019). Water loss over 10% in the medium term is expected to result in a rapid release of water relative to the change in ecosystem water potential (Fig. 2), and the progressive decline of ecosystem function (Grossiord *et al*., 2020). There appears to be a relationship between relative hydraulic capacitance (C_R_) and biomass, such that C_R_ increases by about 1% MPa^−1^ per 100 Mg ha^−1^ biomass (Fig. 6a), although this is only marginally non-significant in the analysis presented here (*P* = 0.091). The mean +/-standard deviation C_R_ is 6.8 +/-2.9 % Mpa^−1^.

**Figure 5.**
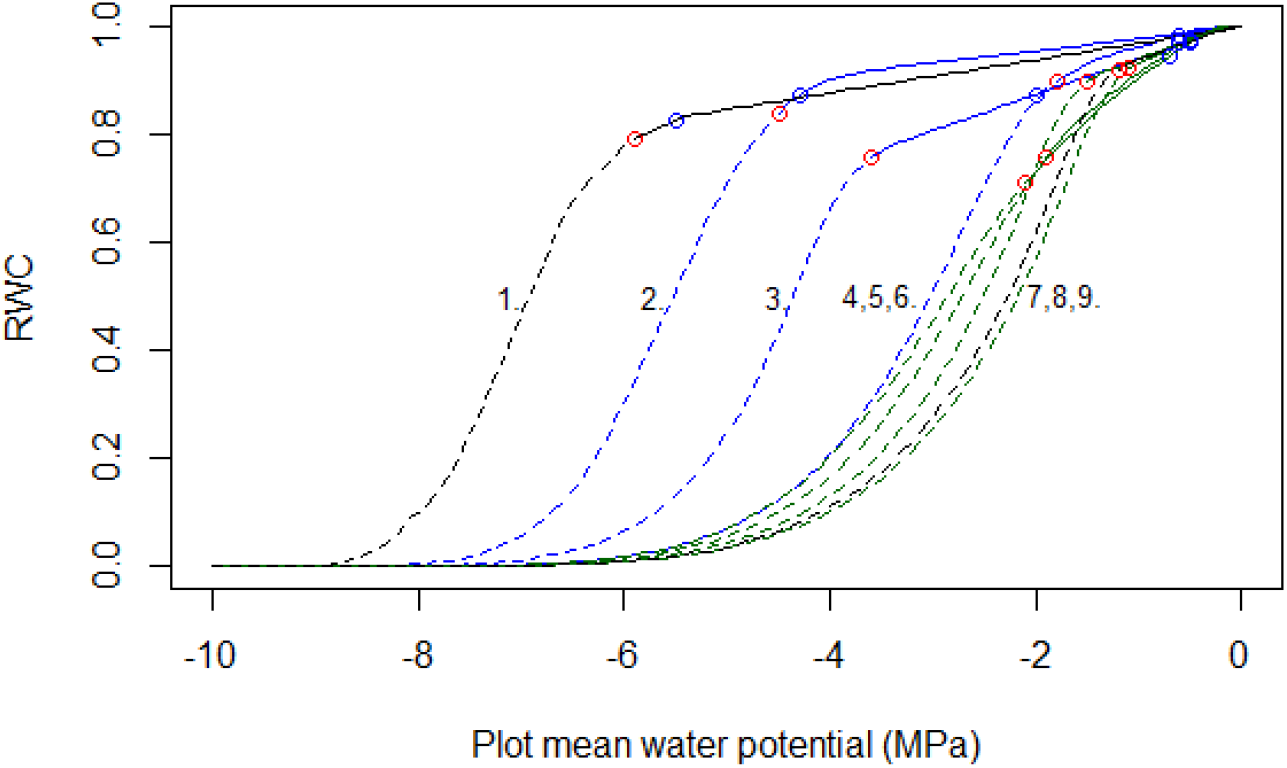
Relative water content versus water potential of the following ecosystems: 1. Alice Mulga; 2. Great Western Woodland; 3. Cumberland Plain; 4. Tumbarumba; 5. Caxiuanã (droughted); 6. Caxiuanã (non-droughted); 7. Cow Bay; 8. Litchfield; 9. Robson Creek (Note: number order differs from other figures for clarity). Solid lines are based on data, while the dashed lines represent approximations of the pressure-volume relationship following the water potential threshold. Tropical rainforest systems in green, temperate forest in blue, and tropical savannah in black. The points on each curve represent equilibrium water potentials as measured at predawn (blue) and the threshold (red) water potentials.

**Figure 6.**
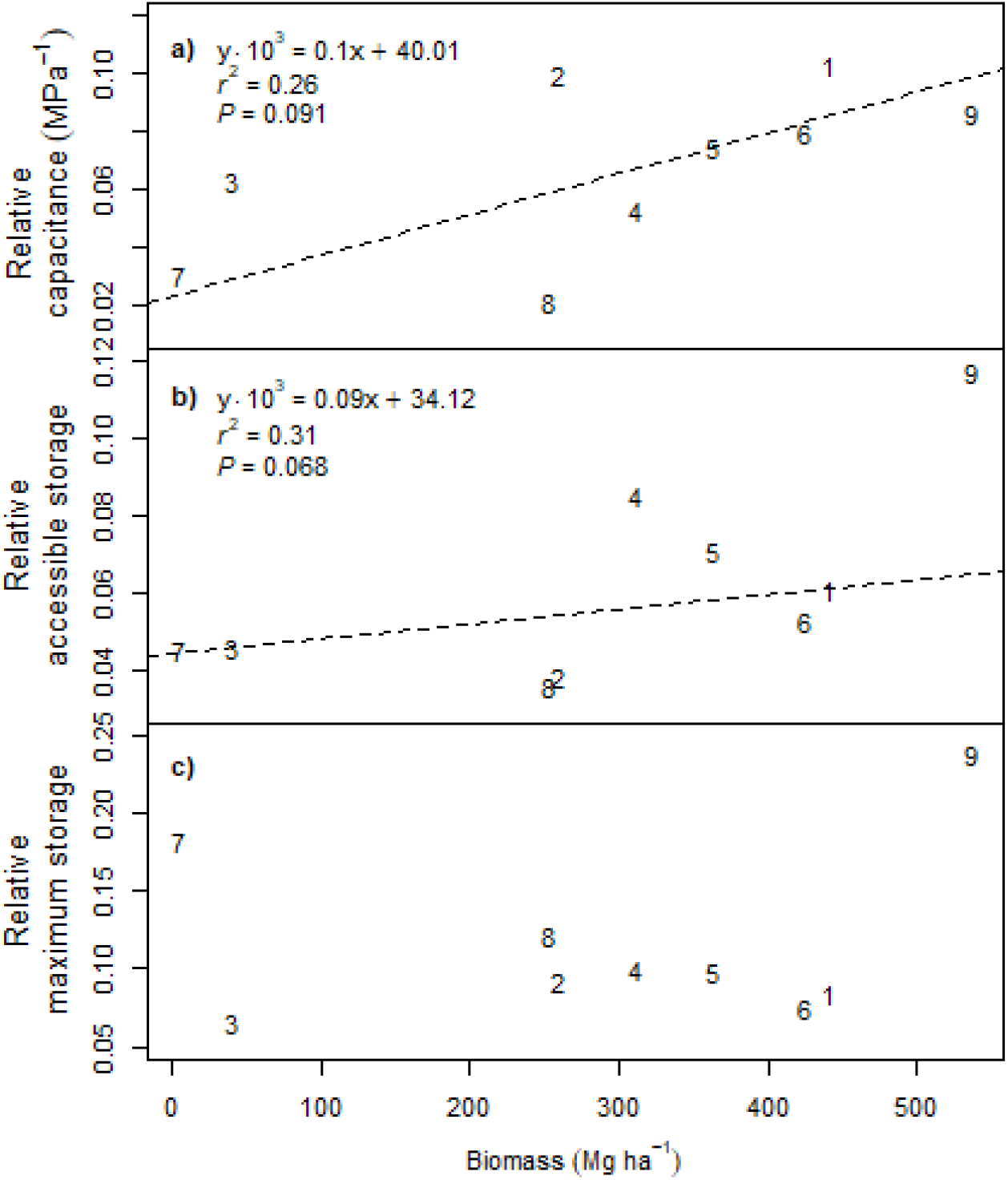
Capacitance (a), accessible (b) and maximum (c) water storage relative to the maximum total water content of each ecosystem. Each number on the plot represents data from each site described in Table 1 corresponding to: 1. Caxiuanã (non-droughted TRF); 2. Caxiuanã (droughted TRF); 3. Litchfield (wet trop. Savanna); 4. Tumbarumba (temp. woodland); 5. Cow Bay (TRF); 6. Robson Creek (TRF); 7. Alice Mulga (semi-arid savanna); 8.Great Western Woodland (temp. woodland); 9. Cumberland Plain (temp. woodland). Regressions in a) and b) are shown as they are nearly significant (*P* < 0.1), but there was no relationship in c) (*P* = 0.737).

The findings that both ecosystem water content and capacitance vary predictably with biomass potentially provide a mechanistic basis for linking remote sensing estimates of landscape water and the ecophysiology of ecosystems.

### 4.3. Limitations and considerations

In this study we only used sapwood water (WC_sw_) in the EPV, where sapwood fraction is possibly the largest source of uncertainty. Quantitatively heartwood water is probably the largest unaccounted for pool, which could influence remote sensing estimates of θ_ES_ particularly in the context of more direct methods, e.g. microwave (Konings *et al*., 2017) and gravity (e.g. ‘GRACE’, (Sheffield *et al*., 2009)). Heartwood water may also play a role in the response of trees to drought through the slow release of water into the sapwood over the medium- to long-term and so is potentially of physiological significance (Guozheng *et al*., 2018, Kalma *et al*., 1998). Bark and leaves likely contain a quantitatively small fraction of the total amount of θ_ES_, but canopy water content in particular will be highly dynamic and contribute disproportionately to the signal from microwave remote sensing estimates (Holtzman *et al*., 2021). Another poorly represented pool is the water contained in trees < 10 cm dbh, which are typically not included in estimates of biomass in forest systems but may influence the change in θ_ES_ in response to prolonged drought. See SI Section 1. for full discussion of water pools.

The analysis was based on changes in equilibrium water potentials, that is, slow changes in water status over the medium term. A transition into drought would initially involve diurnal oscillations in θ_ES_ concurrently with the decline in equilibrium water potential and content. As drought progresses stomatal/canopy conductance declines (Mallick *et al*., 2016, Zeri *et al*., 2014), predawn water potentials decline and the difference between predawn and midday water potentials becomes less (Martínez-Vilalta & Garcia-Forner, 2017); thus, the hydraulic state of the above ground biomass becomes more stable and approaches the equilibrium conditions estimated in this study.

The post-Ψ_threshold_ PV relationship was estimated in this study based on the shape of published sapwood PV curves. For establishing the utility of the notional ecosystem PV curve, the actual shape of the post-Ψ_threshold_ was not important as the parameters used in this analysis were derived from the pre-Ψ_threshold_ part of the curve. What is important to remote sensing applications is whether there is an acute transition point (TP) between the pre- and post-Ψ_threshold_ parts of the curve. The TP in sapwood is typically obvious, and is mechanistically related to the cavitation threshold. As vessels cavitate, they ‘release’ water into the surrounding tissue (Hölttä *et al*., 2009, Tyree & Ewers, 1991), reducing the rate at which water potential declines because the remaining water is distributed amongst a smaller volume of tissue. Whether or not the TP is acute at the ecosystem-level depends on how variable the Ψ_threshold_ is at which the TP occurs between species, individuals and size classes. This is notably demonstrated in figure 2 where the Caxiuanã traces have a gradual TP due to the higher standard deviation of the Ψ_md_ data (based on 161 trees from 36 species (Bittencourt *et al*., 2020). Thus, on the one hand we expect the TP to be harder to detect from RS data in higher diversity communities; but on the other, a smoother TP may result in ecosystems that are less prone to step changes in biomass.

### 4.4. Medium- to long-term changes in ecosystem biomass

In the same way that water is released from cavitation in plant tissues, we propose that a similar process occurs at large spatial and time scales, whereby drought-induced tree mortality results in the available water being redistributed amongst less biomass, maintaining Ψ_ES_ within tolerable physiological limits. This is somewhat evident from the Caxiuanã long-term throughfall exclusion experiment (CTFE) in Amazonian rainforest in Brazil (Fig. 7). The CTFE excluded 50% of the throughfall from 1 ha of rainforest over more than a decade, resulting in elevated mortality and lower biomass (Meir *et al*., 2018). As a result, the drought plot has lower ecosystem water content, while the diurnal range of canopy water potentials remain similar to those in the (non-droughted) control plot. While it is not known whether the drought plot has reached a stable equilibrium value, the experimental result is consistent with the theory presented here, that changes in the hydraulic environment result in co-dependent changes in biomass and θ_ES_ such that the water:biomass ratio is approximately conserved (corresponding to points 1 and 2 in Fig. 4).

**Figure 7.**
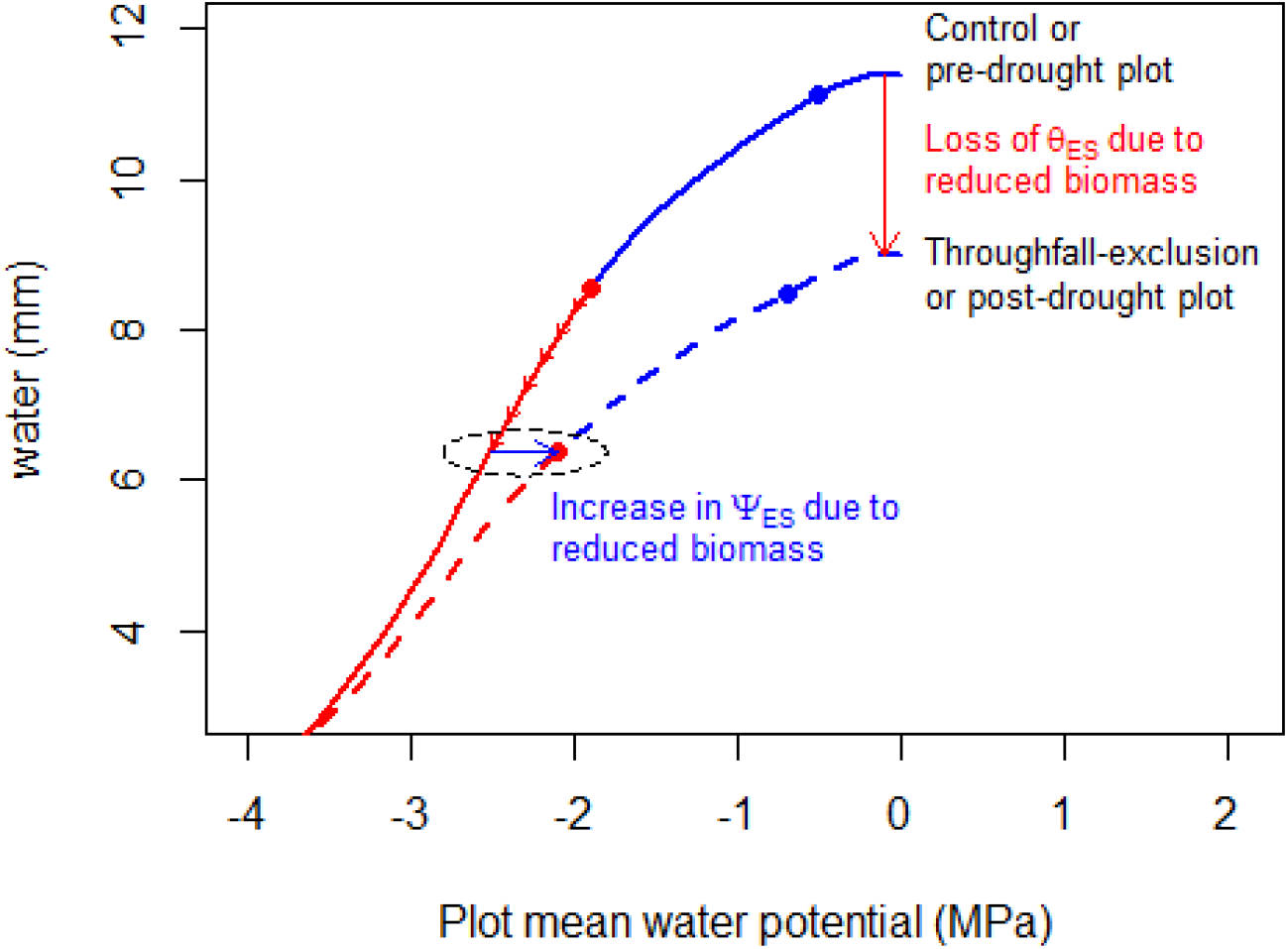
A comparison of the ecosystem water content (θ_ES_) and equilibrium water potential (Ψ_ES_) of the drought plot (thick dashed line) and control plot (thick solid line) in the Caxiuanã throughfall-exclusion experiment, based on data presented in Fig. 2. The red and blue points on each thick line represent the threshold water potential and measured predawn water potentials, respectively. The blue section of each of the thick lines indicates the amount of water available in each plot for ‘reversible’ changes in ecosystem water content, i.e., changes that do not cause physiological damage. The red section of each of the thick lines represent a trajectory of water loss resulting in physiological damage and loss of living tissue/functional biomass. The transition of the drought plot into its current reduced biomass state presumably followed the red arrows on the thick red line to the point at which the biomass reached its current value given the available water. At that point, the equilibrium water potential would have returned to within the normal physiological range represented by the blue arrow (highlighted by the ellipse). The lower threshold water potential in the drought plot (based on measured midday water potentials; Bittencourt et al 2020) might suggest that biomass will continue to decline in the drought plot, given the lack of relationship between midday water potentials and biomass (Fig. S4).

## 5. Conclusion and summary

The concept of the ecosystem pressure-volume curve reconciles our detailed and physically rigorous understanding of small-scale field-measureable processes to the spatial scale applicable to ecosystem and climate science. Successfully bridging that gap in scale potentially allows us to use field measurements to validate remote sensing data, and then remote sensing data to test and update climate models. The ‘state-based’ approach to understanding climate-vegetation feedbacks is based on the principle that ecosystems reach a thermodynamic steady-state with respect to environmental conditions. This assumption allows us to reduce the temporal resolution of data, thereby determining long-term net fluxes of carbon and water from changes in stores, and becoming less dependent on the measurement of processes with high spatial and temporal variability. Acknowledging the existence of additional constraints (e.g. soil nutrients), we propose that to a first approximation the water content of an ecosystem is a direct function of environmental conditions.

We concluded that using the water content of the above ground sapwood, and water potentials during equilibrium (e.g. predawn, or drought) conditions, were practical options for calculating baseline ecosystem PV parameters. This was based on: i) the availability of the existing data, ii) how informative the outcome is in terms of relating to vegetation function, and iii) how useful it could be from the perspective of remote sensing applications. Derivations of both Ψ_ES_ and θ_ES_ could be improved from our estimates with more comprehensive data on water potential, water content and capacitance at larger scale, and better spatial representation across landscapes.

Our first estimates suggest that there appears to be a consistent ratio of ‘physiologically active water’ to biomass across the biomes we looked at of approximately 1:3. Similarly, in absolute terms the water available for reversible changes in θ_ES_, and hydraulic capacitance, also increases with biomass. In relative terms, there were no significant relationships between hydraulic traits and biomass, possibly suggesting these relative values are conserved across ecosystems. Such generalisations across biomes offer the first insight into the utility of the state-based approach for gaining ecophysiologically meaningful interpretations of landscape-scale data.

## Footnotes

1 ‘Equilibrium’ here refers to the thermodynamic concept of a system at maximum entropy, where energy gradients have dissipated and there are no net fluxes. During zero evaporation, the sum of water potential and gravitational potential at any point along the vertical profile is equal to the water potential of soil, i.e., there is no net gradient in the sum of energy potentials. This differs from ‘steady-state’ which refers to a constant gradient and/or constant flux.

