## Supplementary material for "The Ecosystem Pressure-Volume Curve": SI

### The Ecosystem Pressure-Volume Curve
### Supplementary Information

Oliver Binks[1], Patrick Meir[2], Maurizio Mencuccini[1]

1. CREAF, Cerdanyola del Vallès, Barcelona 08193, Spain
2. School of Geosciences, University of Edinburgh EH9 3FF United Kingdom

#### 1. Water stored in above ground biomass - Ecosystem water content, θES

Above ground biomass may be viewed as having four compartments that contain water: sapwood, heartwood, bark and leaves. In this Opinion we base ecosystem water content, $\theta_{ES}$, on sapwood only as the quantitatively dominant, and most physiologically relevant, store of water in above ground biomass.

##### *1.1. Sapwood water content*

Volumetrically, sapwood is likely to contain the majority of ‘physiologically active’ water; that is, water that has a high turnover rate within tissues, and is most closely coupled with diurnal variations in water status. Sapwood water content is also mechanistically related to the hydraulic conductance of woody tissue (Rosner *et al.*, 2019). Therefore, while leaf water content is likely to be more dynamic over the short-term, we expect sapwood water content to dominate the variation of ecosystem water content ($\theta_{ES}$) over the medium-term, e.g., seasonal changes, and periodic droughts.

###### *1.1.1 Deriving the sapwood fraction,* $F_{sw}$

The only study we found providing a direct allometric relationship between diameter at breast height (DBH) and sapwood volume fraction ($F_{sw}$) for whole trees was by Cordero & Kanninen (2003). This study used 87 individuals of a single species (*Tectona grandis*) in Costa Rica. Cordero & Kanninen (2003) derived the following relationship, originally reported in percent and subsequently referred to as C03:

$$F_{sw} = [139.783 - 67.788*\log_{10}(DBH)] / 100 \qquad [C03]$$

However, the trees used in this study had a maximum DBH < 60 cm, and C03 tends to zero at around DBH = 110 cm. Other studies reporting relationships between sapwood area ($A_{sw}$) and tree basal area ($A_T$) or DBH, suggest that sapwood depth increases logarithmically with tree size (Fig. S1) (Aparecido *et al.*, 2016, Aparecido *et al.*, 2019, Horna *et al.*, 2011, Kunert *et al.*, 2017, Moore *et al.*, 2018, Wang *et al.*, 2010).

One possibility for estimating sapwood fraction was to assume a constant ratio between sapwood and heartwood cross-sectional area throughout the whole tree, in which case $F_{sw}$ = $A_{sw}/A_T$; we will subsequently refer to sapwood fraction derived from cross-sectional area as $F_{sw\_a}$. We compared $F_{sw\_a}$ thus derived from the four studies shown in figure S1 with C03 in figure S2. At DBH values less than 40 cm, C03 was quite similar to $F_{sw\_a}$ derived by Kunert et al (2017) and Moore et al. (2018), suggesting that $F_{sw\_a}$ may be a sensible approximation for $F_{sw}$ in some cases at least. We, therefore, derived our own equation ($F_{sw} = 2.9 \cdot DBH^{-0.6}$) which closely matched C03 in the range of DBH = 0-40 cm, but then plateaus at a value of $F_{sw}$ more similar to the values derived from the other studies describing $A_{sw}$ (Fig. S2).

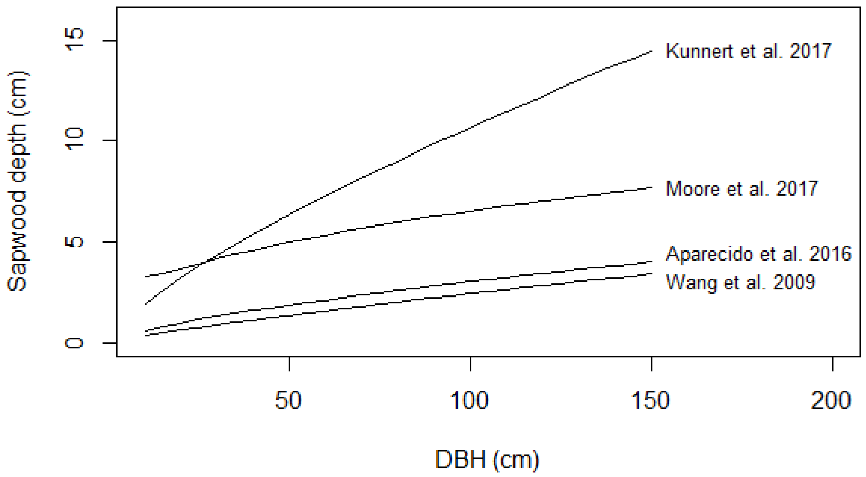

**Figure S1**. Allometric relationships between sapwood depth and tree diameter at breast height (DBH) from four different studies.

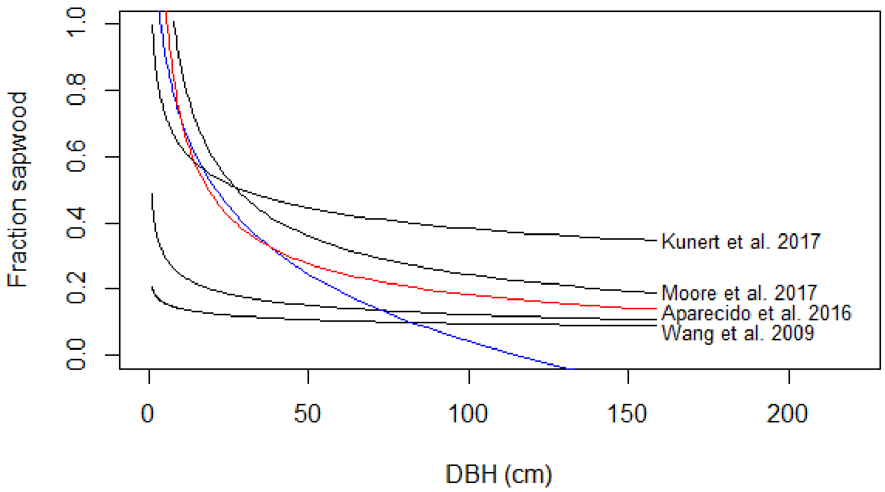

**Figure S2.** Comparison of i) the ratio of sapwood cross sectional area with tree basal area (labelled black lines), with ii) the sapwood volume fraction ($F_{sw}$) relationship derived by Cordero & Kanninen (2003) (C03, blue line), and iii) the modified function derived here to avoid $F_{sw}$ becoming negative (red line).

##### *1.2. Heartwood water content*

Estimates for the water content of heartwood vary widely from 30-220% of the dry mass (Hillis, 1987), and while heartwood is generally considered to be hydraulically isolated from sapwood in terms of diurnal water usage (Beedlow *et al.*, 2007, Cermák *et al.*, 2007, Clark & Gibbs, 1957, Holbrook, 1995), it is plausible (likely, even) that heartwood maintains water potential equilibrium with the sapwood (i.e., its immediate environment) in the medium- to long-term (Guozheng *et al.*, 2018, Kalma *et al.*, 1998). In terms of the total amount of water in above ground biomass, and how it changes over time, heartwood water content is a very large source of uncertainty. Estimates of biomass are based on total woody volume. Our comparisons of $\theta_{ES}$ with biomass effectively assume the heartwood is perfectly dry, which is arguably still a relevant comparison representing the ratio of ecosystem biomass to ‘physiologically active’ water (See ‘Sapwood water content’). However, including the heartwood fraction of water content within the $\theta_{ES}$ term would affect our results in the following ways:

i) The ecosystem-level ratio of water to biomass would increase, making it >1:3 (Fig. 4, main text).

ii) The relative storage fractions would decrease in value, i.e. relative capacitance, relative accessible storage, and relative maximum storage, (Fig. 3a-c, main text).

iii) It may also alter the relationship between $\theta_{ES}$ and AGB from linear to non-linear (figures 3d and 4, main text).

As mentioned in the main text, these considerations are relevant in the context of microwave (Konings *et al.*, 2017) and gravity (e.g. ‘GRACE’, (Sheffield *et al.*, 2009)) remote sensing applications, which provide more direct estimates of $\theta_{ES}$. The possible slow release of water over time from the heartwood to the sapwood during progressive drought also has significant physiological implications to ecosystem resilience, and is a subject that requires further research.

##### *1.3. Bark water content*

There is insufficient data available to estimate the amount of water contained in bark (including phloem) at plot-level with any useful level of certainty. It is probably reasonable to assume that the absolute contribution of bark water to the total water at plot-level is small, perhaps negligible, particularly in high biomass systems. Therefore, we do not address the issue further here, except to acknowledge that bark water content may contribute significantly to buffering short-term fluctuations in water status, and that this is also a topic that requires further investigation.

##### *1.4. Leaf/canopy water content*

Leaves clearly contribute significantly to diurnal oscillations in canopy water content, and canopy water content disproportionately influences the signal of microwave remote sensing applications (Konings *et al.*, 2021, Konings *et al.*, 2017). Moreover, deciduousness and drought-induced reductions in leaf area are mechanisms that result in rapid changes in canopy water content. Therefore, incorporating the dynamics of canopy water relations is important for improving the temporal resolution of estimated ecosystem water content.

In terms of absolute quantities, or relative to the sapwood water content, amount of water contained in leaves is relatively little in high biomass systems. The value of total leaf water content ($WC_{canopy}$, units: mm or kg $m^{-2}$) was calculated for three sites presented in the main analysis (Table S1) as the product of leaf area index (LAI) and leaf water per area (LWA). A mean value of 0.135 kg $m^{-2}{}_{leaf\ area}$ for LWA was calculated using data from Binks et al. (2016b) and Binks et al. (2016a) from 43 trees of 6 genera of Amazonian rainforest trees.

Typically leaf water content at turgor loss point is around 85-90% of the saturated value (Bartlett *et al.*, 2012), suggesting that the contribution of canopy water content to short-term variation in ecosystem water content would only be around 10% of the values in Table S1, assuming no substantial change in leaf area. For perspective, the typical resolution of a rain gauge is around 0.1 mm.

In most woody ecosystems, we expect the loss of the entire canopy leaf area would only change the ecosystem water content by a few percent. The values in Table S1 show total leaf water content as a proportion of sapwood water, as derived in this study. This proportion would be less if heartwood water content was included in the estimate. The fraction of the system detected by microwave remote sensing methods ranges from the including only the upper portion of the canopy to the entire profile of the vegetation including some soil, depending on biomass (Konings *et al.*, 2019). Therefore, the contribution of $WC_{canopy}$ to the signal from microwave remote sensing would be specific to the system.

**Table S1.** Water contained within leaves.

| Site | Leaf area index | Leaf water per ground area (mm) | Leaf water as a percent of plot-level sapwood water content (%) | Data source |
|---|---|---|---|---|
| Daintree Rainforest* | 2.65 | 0.36 | 2.4 | (Yang *et al.*, 2018) |
| Tumbarumba | 1.35 | 0.18 | 3.3 | (Yang *et al.*, 2018) |
| Caxiuana | 5 | 0.68 | 5.9 | (Fisher *et al.*, 2007) |

*Daintree forest is 15 km away from Cow Bay in contiguous forest. The LAI is from Daintree Forest while the ecosystem water content was from Cow Bay.

#### 2. Choosing a threshold water potential, $\Psi_{threshold}$

Our aim here was to choose a threshold water potential i) that delineates a change in performance, e.g., canopy conductance, but ii) occurs before the onset of significant physiological damage, resulting in the potential for rapid recovery. In the absence of exceptional drought stress, the midday water potential of the canopy ($\Psi_{md}$), represents a 'target' water potential, i.e., a physiologically tolerable level of water stress that balances carbon acquisition with hydraulic risk. The water potential at 50 % loss of hydraulic conductance of the stem (the P50), is often used as a reference 'point of no return', but by definition represents a water potential at which physiological function has already been significantly impaired. Therefore, ideally, the threshold water potential would be between the 'safe' water potentials that typically occur at midday and the P50.

Midday canopy water potentials are usually measured in the field under 'typically' stressful conditions, e.g. dry season, when the vegetation is transpiring and there are significant vertical gradients in water potentials. Applying $\Psi_{md}$ as a threshold under equilibrium conditions ($\Psi_{threshold}$) means that the stem water potentials are lower than they would be under typical midday conditions due to the absence of a significant vertical gradient in water potential). However, $\Psi_{md}$ of the canopy is consistently less negative than P50 across many species and vegetation types (Choat *et al.*, 2012). This suggests that a system at water potential equilibrium equal to $\Psi_{md}$ is closer to the point of significant physiological dysfunction than it is under standard midday conditions, but has not necessarily reached a state of significant functional deterioration. Thus using $\Psi_{md}$ as $\Psi_{threshold}$ meets the previously stated requirements.

Another candidate for $\Psi_{threshold}$ is the turgor loss point (TLP), which may well be a suitable alternative for the water potential threshold. However, some plants routinely go below TLP (Bryant *et al.*, 2021) suggesting that it may not represent a limitation to the conductance of water. In practical terms, there also is more data on $\Psi_{md}$ than TLP.

#### 3. Additional figure

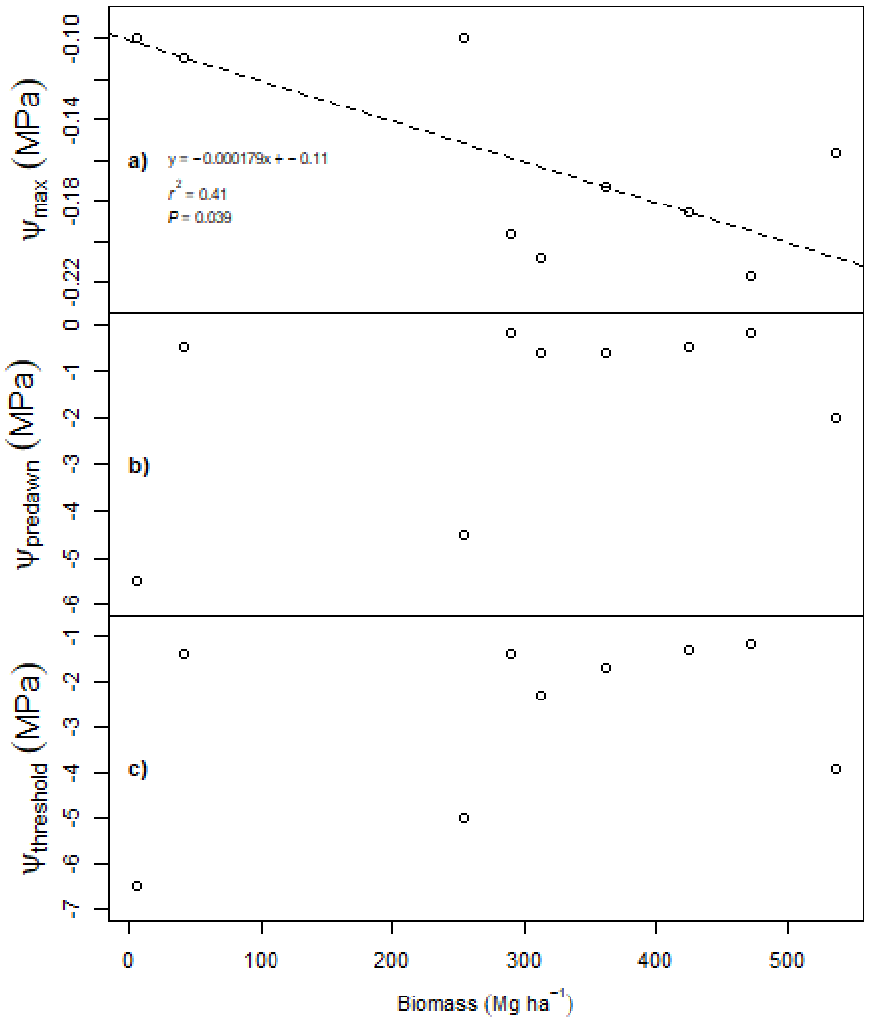

**Figure S4.** Site-level water potentials versus biomass. $\Psi_{max}$ is the maximum possible (least negative) water potential given mean tree height; $\Psi_{predawn}$ is measured canopy predawn water potentials reported from studies in Table 1 in main text; $\Psi_{threshold}$ is the water below which ecosystem-level physiological function begins to decline (here it is the reported dry season midday canopy water potentials).
